# Conserved reward-mediated, reinforcement-learning mechanisms in Pavlovian and instrumental tasks

**DOI:** 10.1101/2022.06.12.495805

**Authors:** Neema Moin Afshar, François Cinotti, David Martin, Mehdi Khamassi, Donna J. Calu, Jane R. Taylor, Stephanie M. Groman

## Abstract

Model-free and model-based computations are argued to distinctly update action values that guide decision-making processes. It is not known, however, if these model-free and model-based reinforcement learning mechanisms recruited in operationally based, instrumental tasks parallel those engaged by Pavlovian based behavioral procedures. Recently, computational work has suggested that individual differences in the attribution of incentive salience to reward predictive cues, i.e., sign- and goal-tracking behaviors, are also governed by variations in model-free and model-based value representations that guide behavior. Moreover, it is not appreciated if these systems that are characterized computationally using model-free and model-based algorithms are conserved across tasks for individual animals. In the current study, we used a within-subject design to assess sign-tracking and goal-tracking behaviors using a Pavlovian conditioned approach task, and, then characterized behavior using an instrumental multi-stage decision-making (MSDM) task in rats. We hypothesized that both Pavlovian and instrumental learning processes may be driven by common reinforcement-learning mechanisms. Our data confirm that sign-tracking behavior was associated with greater reward-mediated, model-free reinforcement learning and that it was also linked to model-free reinforcement learning in the MSDM task. Computational analyses revealed that Pavlovian model-free updating was correlated with model-free reinforcement learning in the MSDM task. These data provide key insights into the computational mechanisms mediating associative learning that could have important implications for normal and abnormal states.

**Significance Statement:** Model-free and model-based computations that guide instrumental, decision-making processes may also be recruited in Pavlovian based behavioral procedures. Here, we used a within-subject design to test the hypothesis that both Pavlovian and instrumental learning processes were driven by common reinforcement-learning mechanisms. Sign-tracking and goal-tracking behaviors were assessed in rats using a Pavlovian conditioned approach task, and, then instrumental behavior characterized using a multi-stage decision-making (MSDM) task. We report that sign-tracking behavior was associated with greater model-free, but not model-based, learning in the MSDM task. These data suggest that Pavlovian and instrumental behaviors are driven by conserved reinforcement-learning mechanisms.

## Introduction

Cues in the environment that predict rewards can acquire incentive value through Pavlovian mechanisms (Flagel et al., 2009) and are necessary for the survival of an organism by facilitating predictions about biologically relevant events that enable an organism to engage in appropriate preparatory behaviors. Pavlovian incentive learning, however, can imbue cues with strong incentive motivational properties that exert control over behavior, which can lead to maladaptive and detrimental behaviors (Saunders and Robinson, 2013). For example, cues that are associated with drug use can enhance craving in addicts, and, because of their control over behavior, may precipitate relapse to drug-taking behaviors in abstinent individuals (Hammersley, 1992). Understanding the biobehavioral mechanisms underlying associative learning could, therefore, provide critical insights into how stimuli gain incentive salience.

Pavlovian associations have largely been presumed to occur through *model-free*, or stimulus-outcome, learning: cues that are predictive of rewards incrementally accrue value through a temporal-difference signal that is likely to be mediated by mesolimbic dopamine (Huys et al., 2014; Nasser et al., 2017; Saunders et al., 2018). Theoretical work has, however, proposed that Pavlovian associations may also involve learning that is described in the computational field as *model-based* (Dayan and Berridge, 2014; Lesaint et al., 2014a) whereby individuals learn internal models of the statistics of action-outcome contingencies. This hypothesis has been supported by data indicating that Pavlovian associations not only represent accrued value, but also the identity of Pavlovian outcomes (Robinson and Berridge, 2013) and by neuroimaging studies that identify neural signatures of model-free and model-based learning in humans during a Pavlovian association task (Wang et al., 2020).

Pavlovian autoshaping procedures have been used to quantify the extent to which animals attribute incentive salience to cues predictive of rewards (Flagel et al., 2009, 2011; Nasser et al., 2015). When animals are presented with a cue associated with food reward delivery, the majority of rats – known as sign-trackers (ST) – will approach and interact with the cue, whereas other rats – known as goal-trackers (GT) – will approach the location of the reward delivery (Hearst and Jenkins, 1974; Boakes, 1977). Rats that display ST behaviors, therefore, attribute incentive salience to the cue, whereas rats that display GT behaviors do not (Uslaner et al., 2006), or at least acquire less incentive to the cue than the goal. Our computational work (Lesaint et al., 2014a; Cinotti et al., 2019) has suggested that these conditioned responses may be linked to individual differences in the extent to which rats use model-free and model-based reinforcement-learning systems to guide their behavior. For example, when using a hybrid reinforcement-learning model to simulate Pavlovian approach behaviors we were able to recapitulate ST behaviors by increasing the weight of model-free updating and, notably, GT behaviors by increasing the weight of model-based updating (Cinotti et al., 2019). Variation in Pavlovian approach behaviors in rodents may, therefore, reflect individual differences in model-free and model-based control over behavior (Dayan and Berridge, 2014; Lesaint et al., 2014a).

Use of the multi-stage decision-making (MSDM) task in humans (Daw et al., 2011; Culbreth et al., 2016) and animals (Miller et al., 2017; Groman et al., 2019a; Akam et al., 2021) has provided empirical evidence that instrumental behavior is influenced by both model-free and model-based reinforcement learning computations. It is not known, however, if the relative contribution of model-free and model-based mechanisms that are recruited in an individual during Pavlovian autoshaping procedures are predictive of their relative contribution during instrumental procedures, such as in the MSDM task (Sebold et al., 2016). If true, this could suggest that the computational mechanisms underlying learning are not unique to Pavlovian or instrumental mechanisms but may represent a common reinforcement-learning framework within the brain that could be useful for restoring the learning mechanisms that are abnormal in disease states (Doñamayor et al., 2021; Groman et al., 2021).

In the current study we sought to test the hypothesis that ST rats would preferentially employ a model-free strategy in an instrumental task, whereas GT rats would preferentially employ a model-based strategy. Pavlovian conditioned approach was assessed in rats (Keefer et al., 2020) before model-free and model-based reinforcement-learning was assessed in a rodent analogue of the MSDM task (Groman et al., 2019a). We report that ST behavior is correlated with individual differences in reward-mediated model-free, but not model-based, learning in the MSDM task. These data suggest that the model-free reinforcement-learning systems recruited during Pavlovian conditioning parallel those recruited in instrumental learning.

## Methods

### Subjects

20 Male Long-Evans rats were purchased through Charles River Laboratories at approximately 6 weeks of age. Rats were pair-housed in a climate-controlled vivarium on a 12hr light/dark cycle (lights on at 7 am; lights off at 7 pm). Rats had ad libitum access to water and underwent dietary restriction to 90% of their free-feeding body weight throughout the experiment. Experimental procedures were approved by the Institutional Animal Care and Use Committee (IACUC) at Yale University and according to the National Institutes of Health institutional guidelines and Public Health Service Policy on humane care and use of laboratory animals.

### Pavlovian conditioned approach

Rats were first trained using a Pavlovian conditioned approach task as previously described (Keefer et al., 2020). During a single trial, a retractable lever (CS) located to the left or right of a food cup was inserted into the chamber for 10s. As the lever retracted, a single sucrose pellet (45 mg; BioServ) was dispensed into the food cup.

This CS-US pairing occurred on a variable-interval 60s schedule and each CS-US pairing was present 25 times in each session. Each rat underwent a single, daily session on the Pavlovian conditioned approach task for five consecutive days. The primary dependent measures collected were latency to approach the lever and food cup as well as the number and probability of interactions rats made with the lever and food cup within each session. These dependent measures were used to generate a Pavlovian score for each session a rat completed (see data analysis). This Pavlovian score is typically referred to as the Pavlovian Conditioned Approach (PCA) score; however, to avoid confusion with the data reduction technique known as principal component analysis (also commonly referred to as a PCA) we refer to this measure as the Pavlovian score to avoid any confusion.

### Deterministic MSDM task

Following the Pavlovian conditioning approach sessions, rats were trained to make operant responses (e.g., nosepokes and lever responses) in order to receive a liquid reward delivery (90 μl of 10% sweetened condensed milk) in different operant environments than those used for the Pavlovian conditioning approach task. Once operant responding had been established, rats were trained on a deterministic MSDM task using procedures previously described (Groman et al., 2019a). In the deterministic MSDM task, choices in the first stage deterministically led to the second stage state.

Second stage choices were probabilistically rewarded. Rats initiated trials by making a response into the illuminated food cup. Two levers located on either side of the food cup were extended into the box and cue lights above the levers illuminated (*s_a_).* A response made on one lever (*s_a_a_1_*) resulted in the illumination of two noseport apertures (e.g., ports 1 and 2, *s_B_*), whereas a response made on the other lever (*s_a_a_2_*) would result in the illumination of two other apertures (ports 3 and 4, *s_C_*). Entries into either of the illuminated apertures were probabilistically reinforced using an alternating block schedule.

Each rat was assigned to one specific lever-port configuration (configuration 1: left lever → port 1,2, right lever → port 3,4; configuration 2: left lever → port 3,4, right lever → port 1,2) that was maintained through the study. Reinforcement probabilities assigned to each port, however, were pseudorandomly designated at the beginning of each session (0.90 vs 0.10 or 0.40 vs 0.15; Figure 4A). Sessions terminated when 300 trials had been completed or 90 min had elapsed whichever occurred first. Trial-by-trial data were collected for individual rats and the probability that rats would select the first stage option leading to the highest reinforced second stage option (p(correct|stage1)) was calculated, as well as the probability that rats would select the highest reinforced second stage option (p(correct|stage2)).

Training on the deterministic MSDM task occurred for three primary reasons: (1) to reduce spatial biases that are common in rats, (2) to ensure rats understood the alternating probabilities of reinforcement at the second stage options, and (3) to verify that rats understood the structure of the task and how first stage choices led to different second-stage options. If rats appreciated the reinforcement probabilities assigned to the second state options and how choices in the first stage influence the availability of second-stage options, then the probability that rats choose the first-stage option leading to the second-stage option with the maximum reward probability (e.g., p(correct | stage 1)) should be significantly greater than that predicted by chance. Rats were trained on the deterministic MSDM until they met the criteria of a p(correct|stage1) being significantly greater than chance in four of the five sessions after completing the 35^th^ training session on the deterministic MSDM. If rats did not meet the criterion after completing 43 sessions on the deterministic MSDM (N=3), training was terminated regardless of p(correct|stage1).

### Probabilistic MSDM task

Choice behavior was then assessed in the probabilistic MSDM task. Initiated trials resulted in the extension of two levers and illumination of cue lights located above each lever. For most trials (70%), first-stage choices led to the illumination of the same second-stage state that were deterministically assigned to that first-stage choice in the deterministic MSDM (referred to as a *common transition*). On a limited number of trials (30%), first-stage choices led to the illumination of the second-stage state most often associated with the other first-stage choice (referred to as a *rare transition*). Second-stage choices were probabilistically reinforced using the same alternating block schedule as that of the deterministic MSDM task. Rats completed 300 trials across five daily sessions on the probabilistic MSDM task.

Trial-by-trial data (∼1500 trials/rat) were collected to conduct logistic regression analyses of decision making (described below). One rat was excluded from all analyses due to an extreme bias in the first-stage choice (e.g., rat chose one lever on 97% of all trials, regardless of previous trial events).

### Data analysis

#### Pavlovian Conditioned Approach

To quantify the degree to which individual rats display ST or GT behaviors, a summary Pavlovian score (historically known as the *Pavlovian Conditioned Approach Index*) was calculated for individual rats by averaging three standardized measures collected in each Pavlovian conditioning session, as previously described (Meyer et al., 2012). These three measures were: 1) a latency score which was the average latency to make a food cup response during the CS, minus the latency to lever press during the CS, divided by the duration of the CS (10 s), 2) a probability score which was the probability that the rat would make a lever press minus the probability that the rat would make a food cup response across the session, and 3) a preference score which was the number of lever contacts during the CS, minus the number of food cup contacts during the CS, divided by the sum of these two measures. This summary Pavlovian score ranged between -1.0 and 1.0, with values closer to 1.0 reflecting a greater prevalence of ST behaviors and values closed to -1.0 reflecting a greater prevalence of GT behaviors. Additionally, a trial-by-trial Pavlovian score was calculated to serve as the dependent measure used in the computational analyses described below. Latency and preference scores were calculated on each trial to generate a trial-specific Pavlovian score. As expected, the average trial-by-trial Pavlovian score was strongly correlated with the summary Pavlovian score (R^2^=0.94; p<0.001).

Previous studies have used the summary Pavlovian score in the last two Pavlovian sessions to classify rats as either exhibiting high or low ST behaviors (Morrison et al., 2015; Rode et al., 2020), as GT behaviors are less commonly observed within the population. We conducted a similar median split of the distribution of summary Pavlovian scores and classified rats as either exhibiting high ST behaviors (N=10) or low ST behavior (N=10). All group-level analyses reported in the current study were conducted using this binary classification.

#### Model-free and model-based learning in the Pavlovian conditioned approach task

We have previously reported that individual differences in Pavlovian approach behavior can be recapitulated using a combination of model-free and model-based reinforcement learning algorithms (Lesaint et al., 2014a; Cinotti et al., 2019). We sought to use this hybrid reinforcement-learning model to index the contribution of both these reinforcement-learning systems to Pavlovian conditioned approach behaviors observed in the current study. This model combines the outputs of these two reinforcement-learning systems to determine on a trial-by-trial basis whether the rat approach either the lever or the magazine. The structure of each trial of the task is represented by a Markovian Decision Process (MDP) consisting of six different states (Figure 1A) defined by the experimental conditions, such as the presence of the lever or of the food, and the current position of the rat (e.g., close to the magazine or the lever). There are five different actions (explore the environment or goE, approach the lever or goL, approach the magazine or goM, engage the closest stimuli or eng_<L>/<M>_, and eat the reward), and state transitions given a selected action are deterministic. For example, if a rat in state 1 (s_1_) chooses the action goL, it will always lead to state 2 (s_2_) whereas if a rat in state 1 (s_1_) chooses the action goM, it will lead to state 3 (s_3_). Action values for all possible actions in the current state are generated by the decision-making model which consists of both a Model-Based (MB) and a Feature Model-Free (FMF) reinforcement-learning algorithm (Figure 1B). The MB and FMF value estimates are combined as a sum into a weighted value determined by the free parameter *ω*. An *ω* parameter closer to 1 indicates that action values are more influenced by the MB computations, whereas an *ω* parameter closer to 0 indicates that action values are more influenced by the FMF computations. The weighted values are fed into a softmax function representing the action selection mechanism.

**Figure 1:**
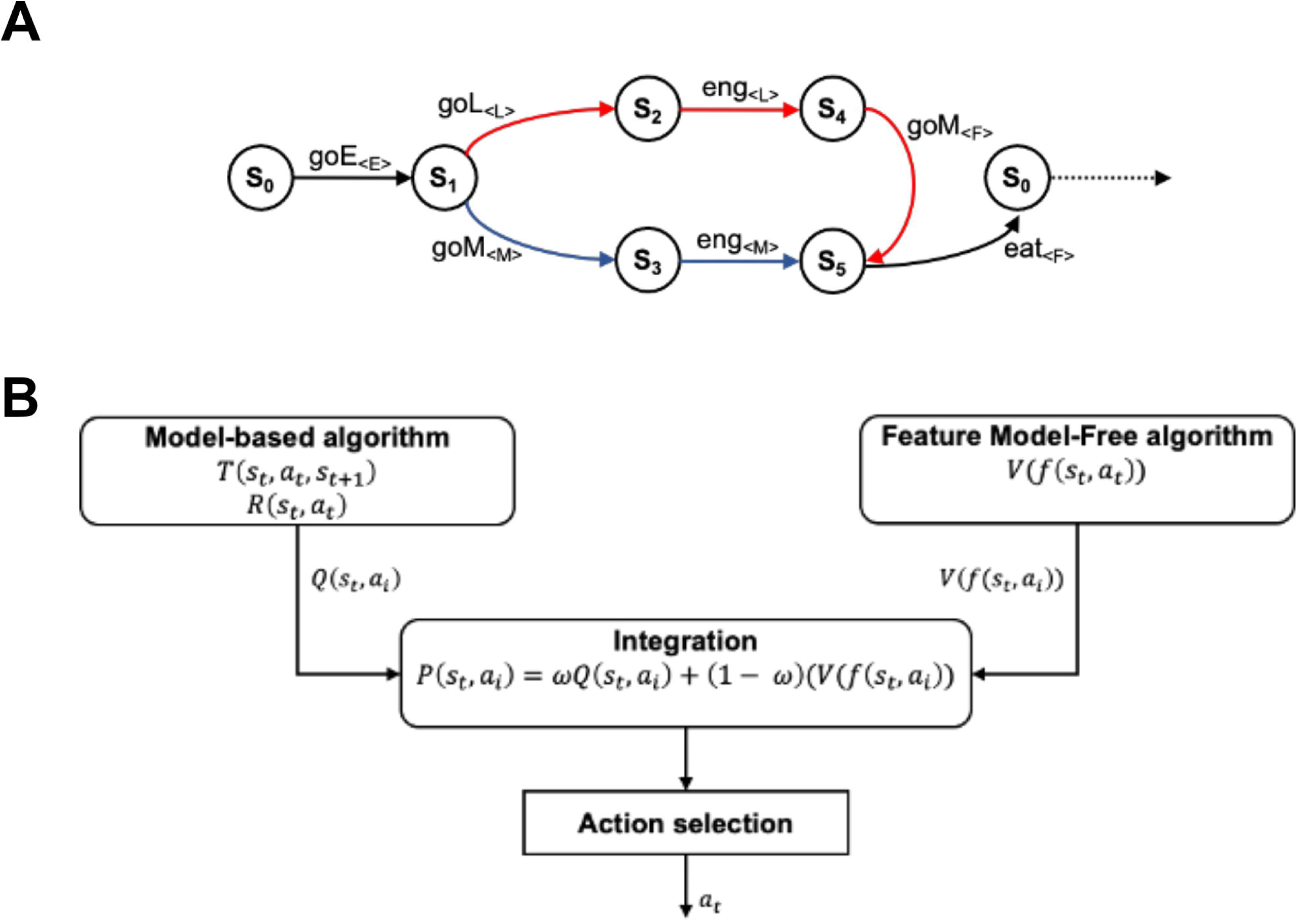
The FMF-MB decision-making model. (A) The Markov decision process of a single trial from the Pavlovian approach task. There are five possible actions leading deterministically from one state to the next: exploring the environment (goE), approaching the lever (goL), approaching the magazine (goM), engaging with the closest stimulus (eng), and eating the reward (eat). Each of these actions focuses on a specific feature indicated in brackets: the environment (E), the lever (L), the magazine (M), and the food (F). These are the features used by the FMF learning component. The red path corresponds to sign-tracking behavior and the blue path to goal-tracking behavior. (B) Schematic of the FMF-MB decision-making model adapted from Lesaint et al. (2014) and Cinotti et al. (2019). The model combines a MB learning system which learns the structure of the MDP and then calculates the relative advantage of each action in a given state, with a FMF system which attributes a value to different features of the environment which is generalized across states (e.g., the same value of the magazine is used in states 1 and 4). The advantage function and value function are weighted by ω, their relative importance determining the sign- vs goal-tracking tendency of the individual and then passed to the action selection mechanism modelled by a softmax function.

The FMF system, compared to instrumental reinforcement-learning algorithms, assigns value representations to the features associated with each action, rather than to the states of the task, which allows a generalization of values between different states. For example, when the rat goes towards the magazine in state 1 (s_1_) or engages the magazine in state 3 (s_3_), it does so motivated by the same feature value (e.g., V(M)) in these two different states which means V(M) is updated twice in the course of a single trial. After each action, the value of the corresponding feature is updated according to a standard temporal difference (TD) learning rule by first computing a reward prediction error (*δ*):

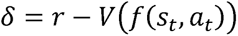

where r is equal to 1 or 0 if reward delivery occurs or not, respectively. This reward prediction error is integrated in the current estimate of the value of the chosen feature

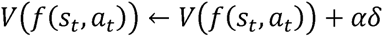

with the learning rate (e.g., *α*) is bounded between 0 and 1. In contrast to our previous FMF algorithm, the discounting parameter *γ* was not included here. This was because our model comparisons (described below in Results) indicated that this parameter was not explaining unique variance in approach behavior of this group of rats.

The TD learning rule was only applied to the selected feature (E, environment; L, lever; M, magazine) in each state transition, except in the case of food, which was equal to 1, the value of reward. Because the rat is likely to visit the magazine during inter-trial interval (ITI), the values of the magazine are revised between state 5 and state 0 according to the following equation:

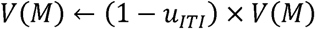

where *u_ITI_* determines the rate at which action values for the magazine (*V*(*M*)) decay during ITI.

The MB system relies on learned transition *T* and reward *R* functions for updating action values. The transition value function aims to determine the probability of going from one state to the next given a certain action. After transitioning from state *s_t_* to *s_t_*_+1_ by performing action at, the transition *T*(*s_t_*,*a_t_*, *s_t_*_+1_) is updated according to the following:

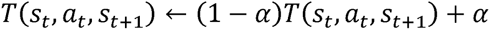

with initial values of *T* set to 0 for all possible state and action combinations. The *T* values for unvisited states are decreased according to the following:

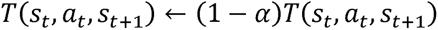

Because the environment is deterministic, *T(s,a,s)* should converge perfectly towards values of 1 for all possible state transitions and remain at a value of 0 for all impossible state transitions (e.g., s_1_➔ s_4_). Similarly, the reward function *R(s,a)* is updated according to the following:

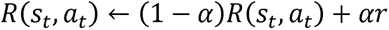

where *r* is set to 1 for (s_5_, eat) and 0 otherwise. Initially, *R(s,a)* is equal to 0 for all state-action pairs. Across actions and experience in each state, *R(s,a)* will converge to a value of 1 for (s_5_, eat) state-action pair and remain at a value of 0 for all other state-action pairs. The action-value functions for each possible action *α_i_* in the current state s*_t_* are then calculated according to the following:

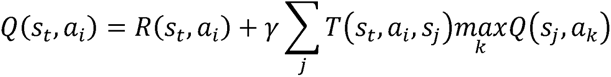

where *β* is the discounting parameter. Once the FMF and MB systems have outputted the feature values and the advantages of the possible actions, these are integrated through a weighted sum:

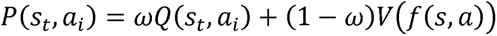

with *ω* bounded between 0 and 1. These integrated values are then entered into a softmax function to compute the probability of selecting each action:

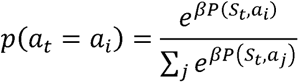

where *β* is the inverse temperature parameter quantifying choice stochasticity.

This model contained five free parameters: the learning rate *α*, the discounting factor *γ*, the inverse temperature *β*, the ITI update factor *u_ITI_*, and the integration factor *ω*. Trial-by-trial behavior was classified as either ST or GT and fit with five free parameters selected to maximize the likelihood of each rat’s sequence of behavior as follows:

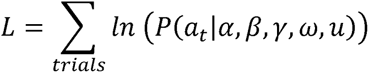

To avoid local maxima, starting values for each free parameter were optimized using a grid search such that each parameter had three possible initial values and all 3^5^ possible combinations were tested as the starting point of the gradient descent procedure to maximize the likelihood *L*. The parameter estimates that yielded the largest log-likelihood were retained and are reported in Table 1.

#### Logistic regression of choice data in the MSDM tasks

We have previously shown that the choice behavior of rats in the MSDM task is guided by previous trial events, such as previous trial outcome, choice, and – in the probabilistic MSDM task – state transitions. Trial-by-trial choice data in the deterministic and probabilistic MSDM was analyzed with a logistic regression model using the glmfit function in MATLAB (MathWorks, Inc. v 2017a). These logistic regression models predicted the likelihood that rats would select the same first-stage choice on the current trial (trial *t*) that they had on the previous trial (trial *t-1*), namely the probability of staying or *p(stay)*. The model used to analyze choice data in the deterministic MSDM contained the following predictors:

Intercept: +1 for all trials, which quantifies the tendency for rats to repeat the same first stage option regardless of any other trial events.

Correct: +1 for trials where the rat selects the first stage option with a common transition leads to the highest reinforced stage 2 option.

-1 for trials where the rat selects the first stage option with a common transition leads to the lowest reinforced stage 2 option.

Outcome: +1 if the previous trial resulted in a rewarded outcome

-1 if the previous trial resulted in an unrewarded outcome

The model used to analyze choice data in the probabilistic MSDM contained the same predictors as described above as well as two additional predictors:

Transition: +1 if the previous trial included a common transition

-1 if the previous trial included a rare transition.

Transition-by-Outcome: +1 if the previous trial included a common transition and was rewarded or if it included a rare transition and was unrewarded.

-1 if the previous trial included a rare transition and was rewarded or included a common transition and was unrewarded.

The “correct” predictor in the logistic regression prevents spurious loading on to the transition-by-outcome interaction predictor (Akam et al., 2015) that can occur when using blocked schedules of reinforcement in the MSDM task. We included the “correct” predictor in all logistic regression models to ensure consistency across analyses and MSDM tasks. Critically, the regression coefficient applied to “outcome” quantifies model-free behavior and the regression coefficient applied to the “transition-by-outcome” interaction quantifies model-based behavior.

### Logistic regression of rewarded and unrewarded outcomes

We found that individual differences in the summary Pavlovian score was related to variation in the outcome regression coefficient (see Results, below). To determine if this relationship was due to differences in the influence of rewarded and/or unrewarded outcomes on choice behavior, we analyzed choice data in the MSDM task using a different logistic regression model that estimated the likelihood that rats would repeat the same first stage choice based on whether the previous trial was rewarded or unrewarded. This logistic regression model, unlike the first, permitted an independent analysis of how each trial outcome (rewarded or unrewarded) influenced first-stage choices. The predictors included in this model were as follows:

Intercept: +1 for all trials. This quantifies the tendency for rats to repeat the same first-stage option regardless of any other trial events.

Rewarded: +1 if the previous trial was rewarded and the rat chose the same lever (first -stage choice) that was selected on the subsequent trial -1 if the previous trial was rewarded and the rat chose a different lever (first-stage choice) than what was selected on the subsequent trial

0 if the previous trial was unrewarded

Unrewarded: +1 if the previous trial was unrewarded and the rat chose the same lever that was selected on the subsequent trial

-1 if the previous trial was unrewarded and the rat chose a different lever than what was selected on the subsequent trial

0 if the previous trial was rewarded

Positive regression coefficients for the rewarded and unrewarded predictor indicate that rats are more likely to persist with the same first-stage choice, whereas negative regression coefficients indicate that rats are more likely to shift their first-stage choice.

### Statistical analyses

Values presented are mean ± SEM, unless otherwise noted. Statistical analyses were conducted in SPSS (version 26; IBM Corp., Armonk NY), MATLAB (version 2017a; Mathworks) and R (https://www.R-project.org). Generalized linear models (GLM; R glmfit package) were used to analyze the relationship between the summary Pavlovian score and choice behavior of rats in the MSDM task. The dependent variable was a binary array coding for whether the first stage choice was the same (+1) or different (0) from the previous trial. Predictors in the model could be correct, outcome, transition, the transition-by-outcome interaction, and summary Pavlovian score or the binary classification of low ST or high ST rats. All higher order (e.g., summary Pavlovian score x outcome x transition) and lower order (e.g., summary Pavlovian score x outcome) interactions were included in the model. Significant interactions were tested using progressively lower order analyses. Another GLM was used to examine the relationship between the summary Pavlovian score and the influence of rewarded and unrewarded outcomes on first-stage choices. The dependent variable was a binary array coding for the first-stage choice (+1 for left lever and 0 for right lever). Predictors in the model were reward, unrewarded, and summary Pavlovian score. All interactions (e.g., summary Pavlovian score x rewarded) were included in the model and significant interactions tested using lower order analyses.

All other analyses were performed in SPSS. Repeated measures data were entered into a generalized estimating equation (GEE) model using a probability distribution based on the known properties of these data. Specifically, event data (e.g., number of trials in which rats chose the highest reinforced first stage option) were analyzed using a binary logistic distribution. Relationships between dependent variables (e.g., *ω* and model-free learning) were tested using the Spearman’s rank correlation coefficient.

## Results

### Pavlovian Conditioned Approach

Pavlovian incentive learning was assessed in rats in a Pavlovian conditioned approach task for five days (Figure 2A,B). The summary Pavlovian score was calculated, and a median split conducted to classify rats as exhibiting either a high (N=10) or low (N=9) ST behaviors (Figure 2C). As expected, the Pavlovian score increased across the sessions in the high ST group (χ^2^ = 91.33; p<0.001) but not in the low ST group (χ^2^ = 0.23; p=0.63; Figure 2D). We then examined how lever and food-cup directed behaviors changed across the five Pavlovian conditioning sessions in both high and low ST rats (Figure 2E-G). Post-hoc analysis of the group (high vs. low ST) x session interaction (χ^2^ = 30.37; p<0.001) indicated that the latency to approach the food cup, the probability of interacting with the lever, and the preference for the lever over the food cup all increased across the Pavlovian sessions in the high ST group (χ^2^ = 68.28; p<0.001), but not in the low ST group (all χ^2^ < 0.99; p>0.32). These session-dependent changes in the high ST rats are similar to observations that we, and others, have reported using Pavlovian conditioned approach tasks (Flagel et al., 2011; Saunders and Robinson, 2011; Keefer et al., 2020).

**Figure 2:**
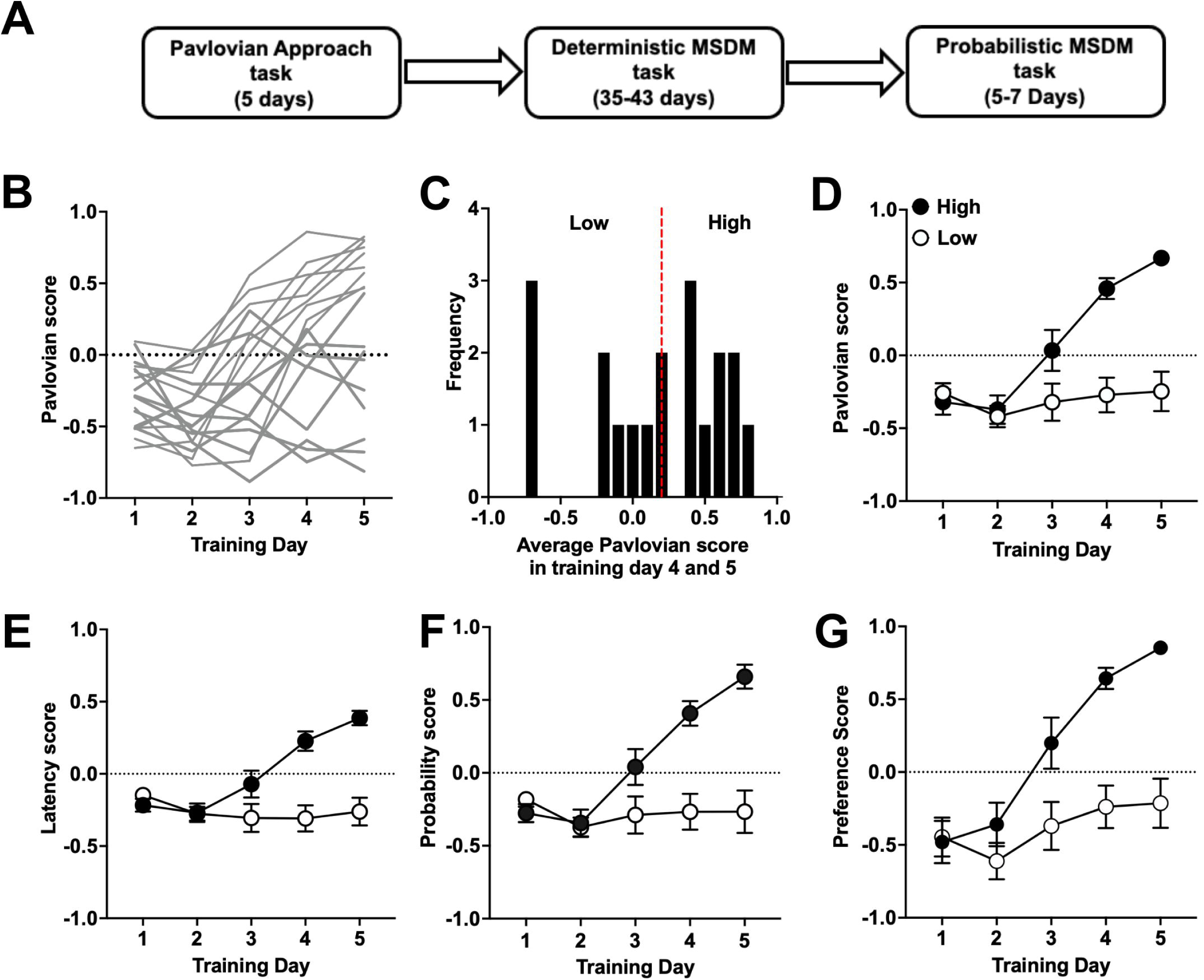
Pavlovian approach task. (A) Schematic of the experimental design. Rats underwent five days of training in the Pavlovian approach task before being trained in the deterministic MSDM task (35-43 days) and tested in the probabilistic MSDM tasks (5-7 days). (B) The Pavlovian score for individual rats across the five days of training. (C) The average Pavlovian score from training day 4 and 5. Rats were divided into two groups based on a median split – low sign-tracking (ST) rats (N=9) and high ST rat (N=10); (D) The Pavlovian score for low ST and high ST rats across the training days. (E) The latency to enter the food cup increased in high ST rats across the training days but did not change in the low ST rats. (F) The probability that rats would interact with the lever increased in the high ST rats across the training days but did not change in the low ST rats. (G) Preference to interact with the lever increased in the high ST rats across the training days but did not change in the low ST rats.

### Computational analysis of Pavlovian approach behavior

Trial-by-trial data from individual rats in the Pavlovian conditioned approach task were fitted with the hybrid model described above and estimates of the five free parameters are presented in Table 1. We also compared the fits of this hybrid model to other variants of this model in which the *ω* was fixed at a value of 1 (e.g., no FMF contribution to the action values) or the *ω* was fixed at a value of 0 (e.g., no MB contribution to the action values). The model in which the *ω* was fixed at a value of 0 had the lowest BIC indicating the FMF-only model best explained the behavior of most rats. This was consistent with the distribution of the Pavlovian scores we observed (see Figure 2C, above) indicating that most rats in the current study exhibited high ST behaviors. The BIC for rats that had the strongest goal-tracking behavior, however, was lowest when the *ω* was fixed at a value of 1, indicating that the full hybrid model is only required for some individuals. These results suggest that although the FMF-only model (e.g., *ω* =0) is sufficient in explaining the behavior of most rats in the current study, this is likely an artifact of the large proportion of ST, and few GT, rats in the current cohort and would not be the case with larger samples sizes consisting of more GT rats.

Because the current study sought to characterize behavioral variation at an individual level, we believe that the hybrid model in which the *ω* is a free parameter is better suited to achieve this goal.

We found that some of the parameter estimates were on extreme ends of the distribution and/or boundary, likely because we were trying to optimize five parameters with a limited number of trials (∼125 trials/rat). In order to improve model fit, we fixed four of the parameters (α,*β*, *γ*, *u_ITI_* ) to the median value estimate obtained from the hybrid model and optimized only the *ω* parameter for each individual rat, as we have previously done (Lesaint et al., 2014a). The BIC of this reduced model was lower than the FMF-only model (Table 2) and the *ω* parameter estimate distribution was found to be less extreme than those observed with the hybrid model (Figure 3A). Nevertheless, the *ω* parameter estimate from the full model was correlated with the *ω* parameter obtained from the restricted model (Spearman’s p=0.87; p<0.001; Figure 3B) suggesting that the restricted model with only a single free parameter (e.g., *ω* parameter) was able to capture the individual differences observed with the full hybrid model.

**Figure 3:**
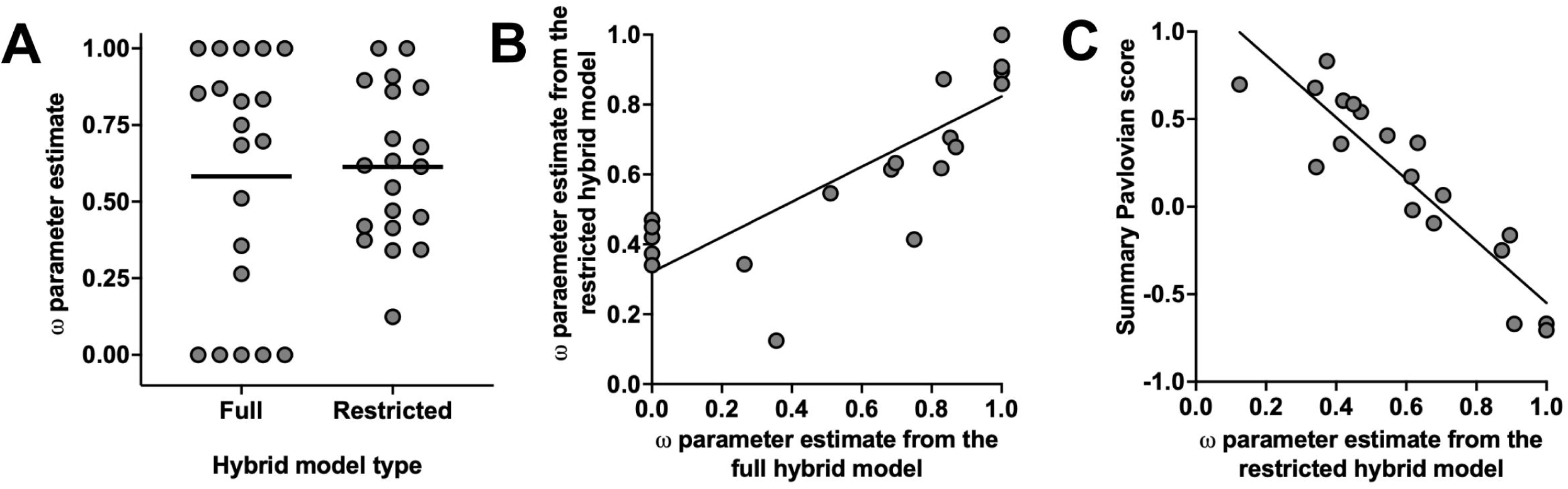
The hybrid reinforcement model for assessing model-based and model-free mechanisms of Pavlovian learning. (A) The *ω* parameter estimate in the full and restricted hybrid model. (B) The relationship between the *ω* parameter in the full hybrid model and the *ω* parameter from the restricted hybrid model. (C) The relationship between the *ω* parameter from the restricted hybrid model and the summary Pavlovian approach score.

Our previous simulation experiments using this hybrid model have found that as the *ω* parameter approaches 0 and the decision-making algorithm favors a FMF system, the prevalence of ST behaviors increases. We hypothesized, therefore, that the *ω* parameter would be negatively correlated with the summary Pavlovian scores across rats. Indeed, individual differences in the *ω* parameter were negatively correlated with variation in summary Pavlovian score (Spearman’s p=0.89; p<0.001; Figure 3C). These results, collectively, indicate that the hybrid FMF reinforcement-learning model can capture meaningful variation in Pavlovian approach behavior.

### Reward-guided behavior in the deterministic MSDM task is related to ST behaviors

Choice behavior on the deterministic MSDM task was then examined (Figure 4A, B). The probability that rats selected the first-stage choice associated with the most frequently reinforced second-stage option increased across the 35 training sessions (*β* = 0.012, p<0.001) and was significantly greater than that predicted by chance in the last five sessions that rats completed (binomial test, p<0.001; Figure 4C). Rats were more likely to repeat a first-stage choice that was subsequently rewarded than a first-stage choice that was subsequently unrewarded (χ^2^ = 113.57, p<0.001; Figure 4D) indicating that second-stage outcomes were able to influence subsequent first-stage choices. These data, collectively, indicate that rats understood the structure of the deterministic MSDM task and, critically, that their first-stage choices influenced the subsequent availability of second-stage options.

**Figure 4:**
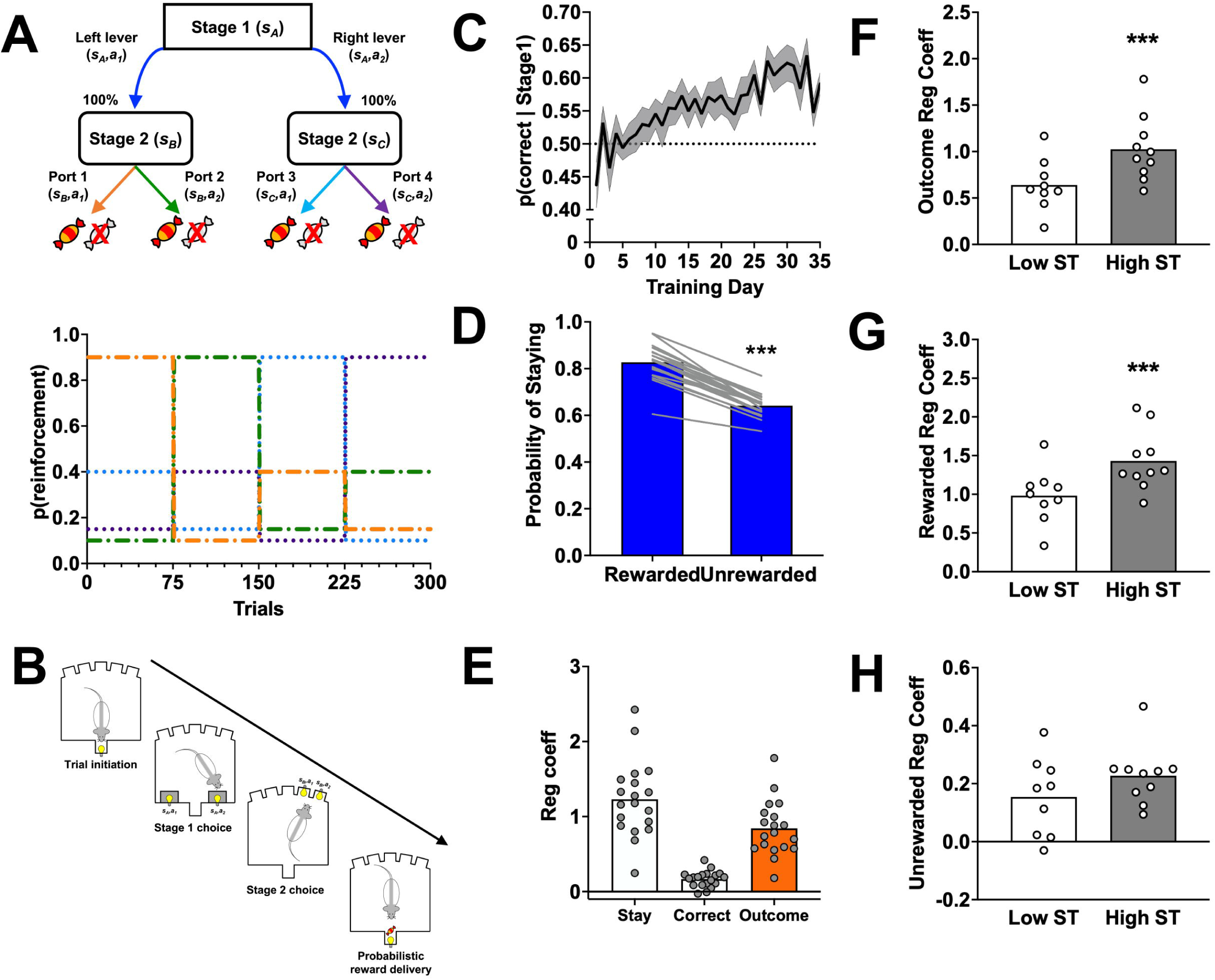
Decision-making in the deterministic MSDM task. (A, top) Rats were trained on the MSDM task in which state transitions were deterministic. (A, below) Stage 2 choices were reinforced according to an alternating block schedule of reinforcement. (B) Schematic of single-trial events. Rats initiated trials by entering an illuminated magazine. Two levers (stage 1) located on either side of the magazine were extended into the operant box and a single lever response led to the illumination of two port apertures (stage 2) located on the panel opposite to the levers. Entries into the illuminate apertures resulted in probabilistic delivery of reward. (C) Probability of selecting the stage 1 option associated with the highest reinforced stage 2 options (p(correct|stage1)) during the first 35 days of training. The probability that choices were at chance level is represented by the dashed line. (D) Probability of choosing the same first-stage choice following a rewarded second-stage choice and following an unrewarded second stage. (E) Regression coefficients for predictors in the logistic regression model from choice behavior of rats in the deterministic MSDM task. (F) Outcome regression coefficient was higher in high ST rats compared to low ST rats. (G) Rewarded regression coefficient was higher in high ST rats compared to low ST rats, but (H) no differences in the unrewarded regression coefficient were observed. Data presented are average ±SEM. *** p<0.001

To quantify the influence of previous trial events (e.g., correct, outcome) on first-stage choices, choice data from rats was analyzed with a logistic regression model (Figure 4E; Table 3). The intercept was significantly greater than 0 (z = 14.92, *p*<0.001), indicating that rats, similar to humans, were more likely to repeat a first-stage choice regardless of previous trial events. Nevertheless, the effect of outcome was also significantly different from 0 (z = 46.56, p<0.001), indicating that rats were using previous trial outcomes (reward and absence of reward) to guide their first-stage choices.

We then examined whether individual differences in Pavlovian approach behavior predicted choice behavior of the same rat in the deterministic MSDM task. The summary Pavlovian score was included as a covariate in the logistic regression model and the two-way interaction between outcome and the Pavlovian score examined. The summary Pavlovian score x outcome interaction was a significant predictor in the model (z=7.51; p<0.001; Table 3) and post-hoc analyses indicated that the regression coefficient for outcome was significantly greater in high ST rats compared to the low ST rats (z=9.58; p<0.001; Figure 4F). These data demonstrate that high ST rats were more likely to use previous trial outcomes to guide their choice behavior compared low ST rats.

The outcome regression coefficient quantifies the degree to which both rewarded and unrewarded outcomes guide subsequent choice behavior. Differences in the outcome regression coefficient that we observed between high and low ST rats might, therefore, reflect variation in how these rats use rewarded or unrewarded outcomes to guide their behavior. To independently assess the impact of rewarded and unrewarded trials on first-stage choices, we conducted a second logistic regression analysis of choice data in the deterministic MSDM task. The rewarded regression coefficient was positive (*β*=1.98 ± 0.03; z=71.59; p<0.001) indicating that rats repeated first-stage choices that resulted in reward. The unrewarded regression coefficient was also positive (*β*=0.33 ± 0.02; z=17.78; p<0.001), but smaller than that for rewarded regression coefficient (χ^2^=106; p<0.001), indicating that rats were more likely to repeat rewarded first-stage choices than unrewarded first-stage choices.

We then examined if the summary Pavlovian score interacted with the rewarded or unrewarded regression coefficients to predict first-stage choices in the deterministic MSDM (Table 4). The interaction between the summary Pavlovian score x rewarded regression coefficient was significant (z=8.93; p<0.001; *β* =0.51) and post-hoc analyses between the low and high ST groups indicated that the rewarded regression coefficient was greater in high ST rats compared to low ST rats (z=12.89; p=0.001; Figure 4G).

The summary Pavlovian score x unrewarded interaction, however, was not significant (z=0.27; p=0.79; *β* =0.01; Figure 4H). High ST rats, therefore, used rewarded outcomes to guide their first-stage choices to a greater degree than low ST rats suggesting that these former individual differences in Pavlovian incentive learning are associated with variation in reward-guided instrumental behavior.

### Probabilistic MSDM task and relationship to Pavlovian approach behavior

To determine if the relationship between the summary Pavlovian score and reward-guided behavior in the above deterministic version of the MSDM task was associated specifically with model-free or model-based reinforcement learning, the choice behavior of rats was assessed in the probabilistic version of the MSDM task (Figure 5A). According to model-free theories of reinforcement learning, the probability of repeating a first-stage choice should be influenced only by the previous trial outcome, regardless of whether the state transition was common or rare (Figure 5B, left). In contrast, model-based theories of reinforcement learning posit that the outcome at the second stage should affect the choice of the first-stage option differently based on the state transition that was experienced (Figure 5B, right). Evidence in humans and in our previous rodent studies, however, indicates that individuals use a mixture of model-free and model-based strategies in the probabilistic MSDM task. Indeed, the probability that rats in the current study would repeat the same first-stage choice according to outcomes received (rewarded or unrewarded) and the state transitions experienced (common or rare) during the immediately preceding trial indicated that rats were using both model-free and model-based learning to guide their choice behavior (Figure 5C).

**Figure 5:**
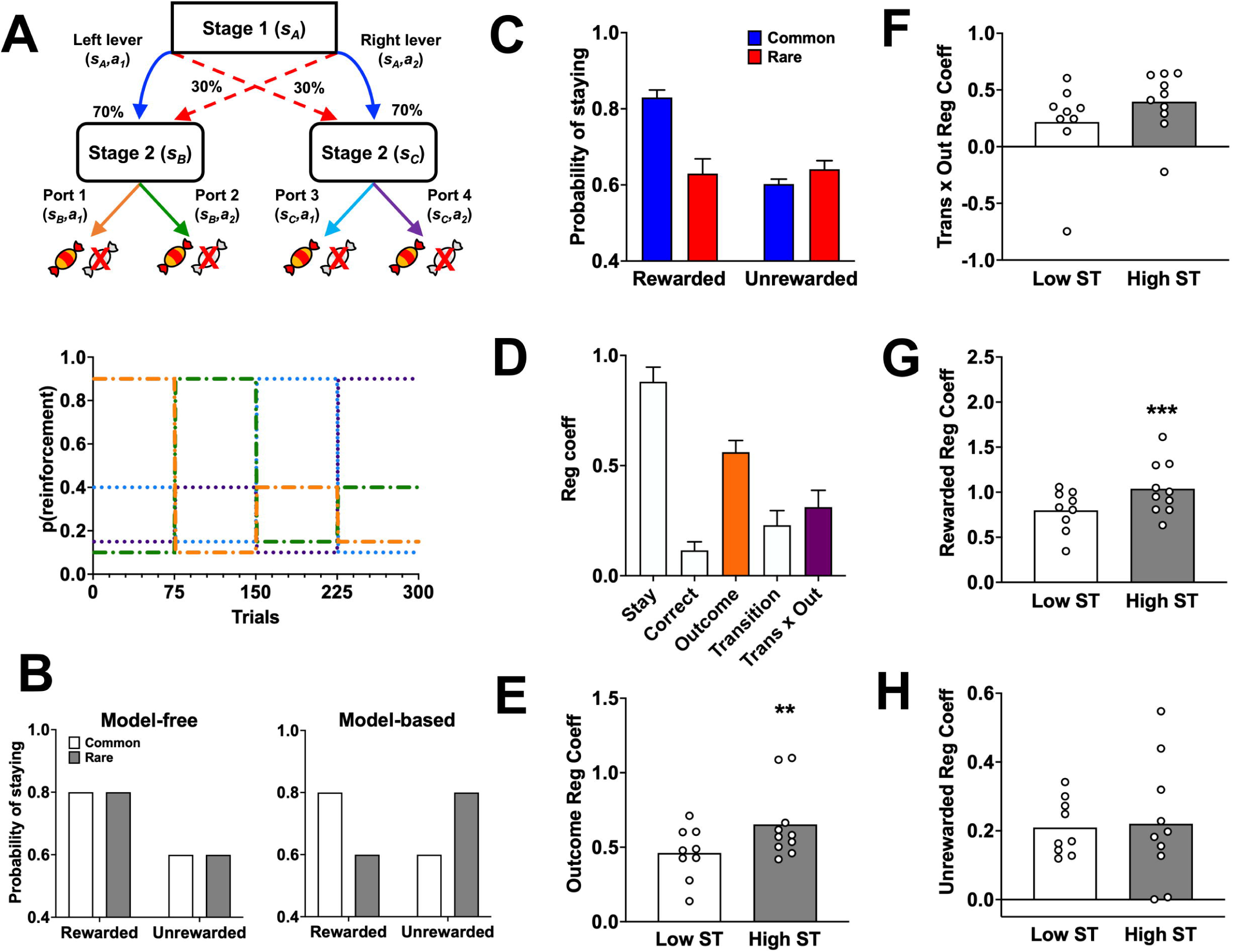
Decision-making in the probabilistic MSDM task. (A) Choice behavior was assessed in the probabilistic MSDM task, which was similar in structure to the reduced MSDM, but state transitions were probabilistic. (B) Probability of staying with the same first-stage choice based on the previous trial outcome (rewarded vs unrewarded) and the state transition (common: open bars; rare: gray bars) in hypothetical data for a pure model-free agent, a pure model-based agent. (C) Probability of staying with the same first-stage choice based on the previous trial outcome (rewarded vs unrewarded) and the state transition (common: blue bars; rare: red bars) during the probabilistic MSDM task reinforced using the alternating schedule. (D) Regression coefficients for predictors in the logistic regression model from choice behavior of rats in the probabilistic MSDM reinforced using the alternating schedule. The coefficient for the outcome predictor (orange bar) represents the strength of model-free learning, whereas the transition-by-outcome interaction predictor (purple bar) represents the strength of model-based learning. (E) The outcome regression coefficient (e.g., model-free learning) was greater in high ST rats when compared to low ST rats. (F) The transition-by-outcome regression coefficient did not differ between low ST and high ST rats. (G) The rewarded regression coefficient was greater in high ST rats compared to low ST rats. (H) The unrewarded regression coefficient did not differ between low and high ST rats. ** p<0.01; *** p<0.001.

To quantify the influence of model-free and model-based strategies, choice data was analyzed with a logistic regression model (Akam et al.,(Daw et al., 2011; Akam et al., 2021) 2015; Groman et al., 2019a, 2019b). The main effect of outcome, which provides an index of model-free learning, was significantly greater than zero (z = 22.65, *p*<0.001; Figure 5D, orange bar) indicating that rats were using second-stage outcomes to guide their first-stage choices. The interaction between the previous trial outcome and state transition, which provides an index of model-based learning, was also significantly greater than zero (z = 15.38, p<0.001; Figure 5D, purple bar). The combination of a significant main effect for outcome and a significant transition-by-outcome interaction suggests that rats were using both model-free and model-based strategies to guide their choice behavior in the probabilistic MSDM task.

We then examined whether the summary Pavlovian score interacted with model-free and/or model-based learning to predict the probability of repeating the same first-stage choice in the probabilistic MSDM task (Table 5). The interaction between the summary Pavlovian score and trial outcome significantly predicted choice behavior (z=3.16; p=0.002), but the interaction between the summary Pavlovian score and the outcome-by-transition predictor did not (z=1.60; p=0.11). Post-hoc comparisons between low and high ST rats indicated that the outcome regression coefficient was significantly greater in high ST rats compared to low ST rats (z=2.67; p=0.008; *β*=0.09; Figure 5E), which was a similar effect observed in the deterministic task (see Figure 4F, above). The outcome-by-transition regression coefficient did not differ between the low and high ST rats (Figure 5F). These data, collectively, suggest that high ST rats rely to a great degree on model-free learning in the MSDM task compared to low ST rats.

To determine if the summary Pavlovian score was associated with rewarded or unrewarded outcomes, choice behavior in the probabilistic MSDM task was analyzed with an alternative logistic regression model. Similar to what we had observed in the deterministic MSDM task, the interaction between the summary Pavlovian score and rewarded predictor was significant (z=4.31; p<0.001; *β* =0.23): high ST rats were more likely to repeat a first-stage choice that led to a rewarded second-stage choice compared to low ST rats (Figure 5G). We also observed a significant interaction between the summary Pavlovian score and the unrewarded predictor (z=-2.69; p=0.007; *β* =–0.09), but the unrewarded regression coefficient was not statistically different between low and high ST rats (Figure 5H). These results, collectively, indicate that individual differences in Pavlovian approach behavior are associated with variation in reward-mediated, model-free learning.

### Pavlovian ST behavior is associated with reward-based, model-free updating

We found that Pavlovian conditioned approach behaviors were associated with reward-mediated, model-free learning in both the deterministic and probabilistic MSDM tasks. This suggests that the model-free computations that guide Pavlovian approach behaviors (e.g., FMF learning) may be related to the model-free computations that influence operant choice behavior in the MSDM task. To test this directly, we compared the regression coefficients obtained from the MSDM task in rats who either had a small *ω* (e.g., more model-free updating in the Pavlovian conditioned approach task) or large *ω* (e.g., more model-based updating in the Pavlovian conditioned approach task) parameter estimate (Figure 6). We hypothesized that if the Pavlovian FMF mechanisms were related to the operant-based model-free learning then the outcome regression coefficient from the MSDM task would differ in rats with a smaller *ω* parameter estimate (e.g., greater FMF updating) compared to rats with a large *ω* parameter estimate (e.g., greater MB updating). As predicted, the outcome regression coefficient (e.g., model-free learning) was larger in rats with a smaller *ω* parameter compared to rats with a large *ω* parameter (χ^2^=6.22; p=0.01; Figure 6A). These differences were specific to model-free learning, as the outcome-by-transition regression coefficient – a measure of model-based learning – did not differ as a function of the *ω* parameter (χ^2^=1.21; p=0.27; Figure 6B). Furthermore, when we compared the rewarded and unrewarded regression coefficients between rats with either a high or low *ω* parameter, only the rewarded regression coefficient differed between the groups (rewarded: χ^2^=6.51, p=0.01, Figure 6C; unrewarded: χ^2^=1.42, p=0.23, Figure 6D). These data suggest that the model-free reinforcement-learning systems recruited during Pavlovian conditioning parallel those recruited in the instrumental MSDM task.

**Figure 6:**
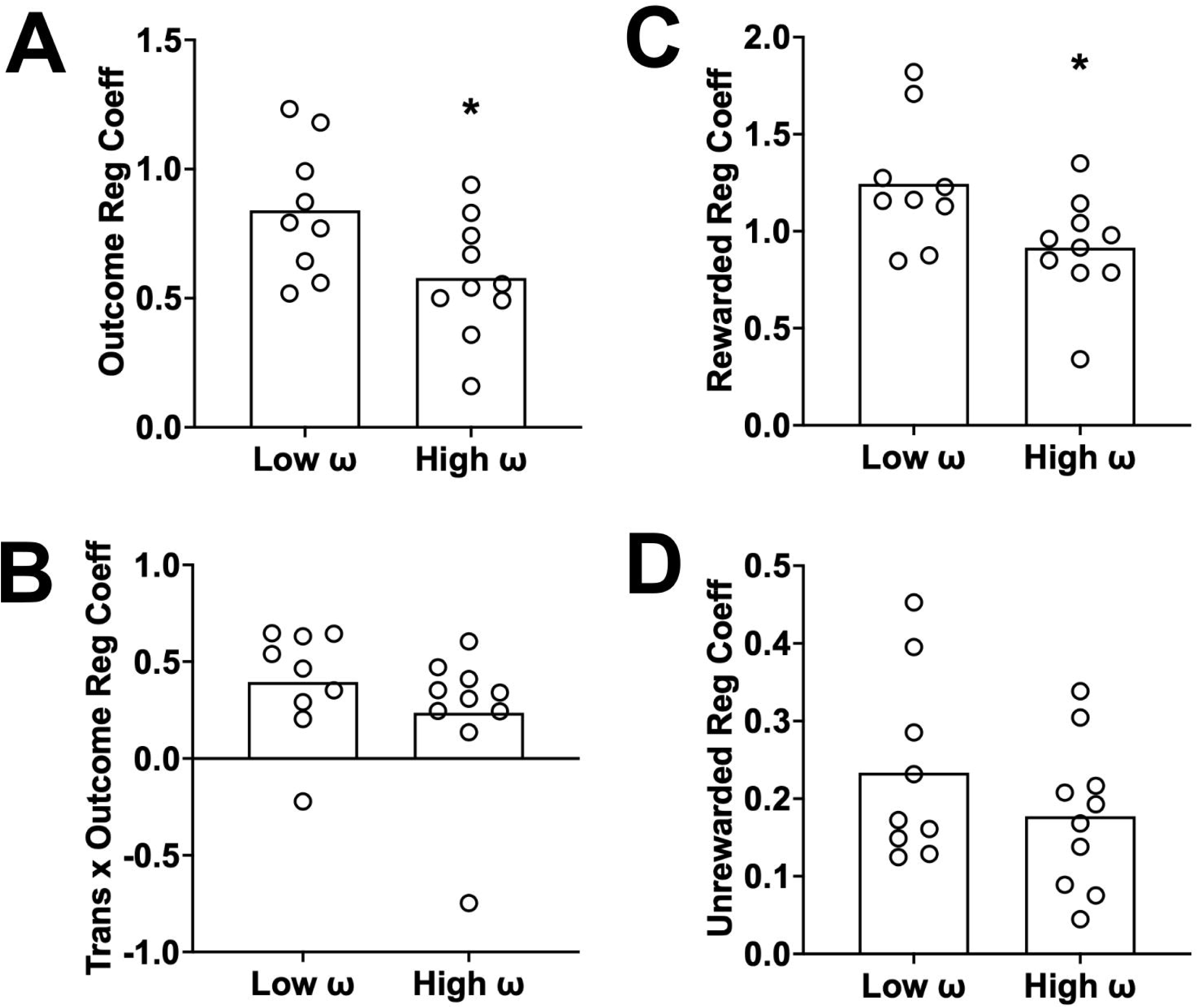
Model-free learning systems in the MSDM are related to model-free learning systems in the Pavlovian approach task. Behavior in the Pavlovian approach task was quantified with the hybrid reinforcement learning model and the degree to which rats were using model-based and/or feature model-free learning to guide their behavior assessed with the free parameter *ω*. (A) Choice behavior of rats with a higher *ω* parameter in the MSDM task was less influenced by model-free learning (e.g., outcome regression coefficient) compared to rats with a lower *ω* parameter. (B) These differences were not observed in model-based learning, as the transition-by-outcome regression coefficient did not differ between rats with a high and low *ω* parameter. (C) Choice behavior of rats with a high *ω* parameter in the MSDM was less influence by rewarded outcomes compared to rats with a low *ω* parameter. (D) The influence of unrewarded outcomes on choice behavior of rats in the MSDM task did not differ between rats with a high or low *ω* parameter.

## Discussion

The current study provides new evidence that the model-free mechanisms that are utilized during the Pavlovian conditioned approach task are related to the model-free mechanisms that guide instrumental decision-making behaviors. We report that a greater prevalence of ST behaviors in the Pavlovian approach task is associated with greater model-free, but not model-based, learning in the MSDM task. Differences in model-free updating observed in high and low ST rats were associated specifically with reward-guided behaviors: rats with higher ST behaviors were more likely to repeat a rewarded choice than rats with lower ST behaviors. No differences in choice behavior following an unrewarded outcome were observed between low and high ST rats. Our data, collectively, provide direct evidence indicating that individual differences in ST behaviors are associated with reward-based, model-free computations. These results suggest that the model-free mechanisms mediating Pavlovian approach behaviors are controlled by the same model-free computations that guide instrumental behaviors and may utilize conserved learning systems that are known to be altered in psychiatric disorders.

### Individual differences in model-free computations are conserved across instrumental and Pavlovian tasks

Rats with higher ST behaviors in the Pavlovian approach task were found to have greater model-free reinforcement-learning in both the deterministic and probabilistic MSDM tasks. These data suggest that the mechanisms that assign and update incentive value to cues predictive of rewards might be the same as those that update representations following rewarded actions. The logistic regression analyses of choice behavior in the MSDM task indicated that rats with higher ST behaviors were more likely to repeat rewarded actions compared to rats with lower ST behaviors. This suggests that the degree of action value updating following rewards was greater in rats with higher ST behaviors and may explain why rats with greater ST behaviors are more resistant to outcome devaluation and slower to extinguish to reward-predictive cues compared to GT, or lower ST, rats (Morrison et al., 2015; Nasser et al., 2015; Ahrens et al., 2016; Smedley and Smith, 2018; Fitzpatrick et al., 2019; Amaya et al., 2020; Keefer et al., 2020). For example, cached representations of actions or cues predictive of rewards may be exaggerated in individuals with greater ST behaviors and, consequently, slower to adjust to changes in the value of the outcome. This may explain why ST behaviors in rats are associated with suboptimal choice behavior in a gambling task (Swintosky et al., 2021).

We did not, however, observe a relationship between Pavlovian approach behaviors and model-based updating in the MSDM task. This was surprising given our previous theoretical work and the experimental work of others (Lesaint et al., 2014b; Cinotti et al., 2019). The lack of association between the Pavlovian summary score and model-based learning in the MSDM task is likely because we only observed a limited number of GT rats in the current sample. Specifically, only three rats in the current cohort of twenty would have been classified as GT rats (see Figure 2, above). This was not because the distribution of Pavlovian approach behaviors in the current study was abnormal – previous studies using larger sample sizes than the current study (e.g., N=560 vs. N=20) have observed similarly skewed distributions (Fitzpatrick et al., 2013).

Future studies that employ large sample sizes to obtain behavioral measures which span the distribution of Pavlovian approach behaviors may, therefore, find a relationship between goal-directed behaviors and model-based learning.

### Neurobiological mechanisms

Although the neurobiological mechanisms underlying Pavlovian and instrumental learning are not fully understood, dopamine neurotransmission is likely to be a point of convergence between ST behaviors and reward-guided, model-free updating. Midbrain dopamine neurons are known to encode reward-prediction errors (RPEs), which is a fundamental computation in model-free learning (Hollerman and Schultz, 1998). The results of studies using voltammetry to quantifying changes in dopamine concentration in the nucleus accumbens – a main output of midbrain dopamine neurons – have proposed that phasic dopamine signals in ST rats is how incentive salience is transferred from the outcome to cue(s) predictive of reward (e.g., lever extension; Flagel et al., 2011). These dopaminergic RPEs were not observed in goal-tracking rats suggesting that variation in attribution of incentive salience may reflect underlying differences in dopaminergic RPEs (Derman et al., 2018; Lee et al., 2018). Indeed, antagonism of dopamine signaling in the nucleus accumbens attenuates the expression of ST behaviors (Saunders and Robinson, 2012).

Dopamine, however, has also been implicated in model-based reinforcement learning. Individual differences in [18F]DOPA accumulation and dopamine tone in the nucleus accumbens of humans and rats, respectively, are associated with variation in model-based learning in the MSDM task (Deserno et al., 2015; Groman et al., 2019a).

Dopamine may play a role in both reinforcement-learning systems. Indeed, recent studies have reported that both model-free and model-based calculations are encoded in the activity of midbrain dopamine neurons (Sadacca et al., 2017; Sharpe et al., 2017; Keiflin et al., 2019), but the influence of these dopaminergic neurons over behavior – and likely learning systems – is mediated by functionally heterogeneous circuits (Keiflin and Janak, 2015; Saunders et al., 2018). For example, mesocortical dopaminergic projections may encode model-based computations whereas mesostriatal/mesopallidal dopaminergic projections may encode model-free computations (Chang et al., 2015).

Studies that integrate circuit-based imaging approaches with biosensor technology (e.g., DLIGHT) to measure circuit-specific dopamine transients in behaving animals could help resolve these critical questions regarding the functional role of dopamine circuits in these learning mechanisms (Kuhn et al., 2018).

### Implications for addiction

Differences in the degree to which individuals attribute incentive salience to cues predictive of reward have been hypothesized to confer vulnerability to addiction. Indeed, there is evidence that ST rats will work hard to obtain cocaine (Saunders and Robinson, 2011), show greater cue-induced reinstatement (Saunders and Robinson, 2010; Saunders et al., 2013; Everett et al., 2020), are resistant to punished drug use (Saunders et al., 2013; Pohořalá et al., 2021), have a greater propensity for psychomotor sensitization (Flagel et al., 2008), and, also display a higher preference for cocaine over food (Tunstall and Kearns, 2015) compared to GT rats. Drug self-administration in short access sessions, however, does not differ between ST and GT rats (Saunders and Robinson, 2011; Pohořalá et al., 2021). These data suggest that drug reinforcement may be similar between ST and GT rats, but that ST rats may be more susceptible or prone to developing compulsive-like behaviors following initiation of drug use.

Only a few studies have used the MSDM task to examine the role of model-free and model-based learning in addiction susceptibility. In a recent study we reported that individual differences in model-free learning in the MSDM task were predictive of methamphetamine self-administration in long-access sessions (Groman et al., 2019b). This relationship, however, was negative: rats with lower model-free learning in the MSDM task took more methamphetamine than rats with higher model-free learning.

Although additional addiction-relevant behaviors were not assessed in this previous study (e.g., progressive ratio, extinction, or reinstatement), the negative relationship between model-free learning and methamphetamine self-administration is surprising given the positive relationship between model-free learning and ST behaviors we observed here. These data might suggest a dynamic role of model-free learning in the different stages of addiction susceptibility (Kawa et al., 2016). For example, greater model-free learning prior to drug use may protect against drug intake but render individuals more vulnerable to the detrimental effects of the drug when ingested.

Indeed, ST rats are less sensitive to the acute locomotor effects of cocaine but have a greater propensity for psychomotor sensitization (Flagel et al., 2008). Future studies that assess Pavlovian conditioned approach behaviors and instrumental reinforcement-learning mechanisms in the same individual prior to evaluating drug-taking and –seeking behaviors may provide a greater understand of the biobehavioral mechanisms underlying addiction susceptibility.

### Summary

The current manuscript provides direct evidence linking incentive salience processes with reward-guided, instrumental behaviors in adult male rats. Our data suggest that Pavlovian approach behaviors and choice behavior of rats in a multi-stage decision-making task are driven by conserved model-free reinforcement-learning mechanisms that are known to be altered in individuals with mental illness, such as addiction (Groman et al., 2022). Determination of whether augmented model-free processes are linked to addiction vulnerability or to the consequences drug-induced circuitry alterations both could be new avenues of translational-based computational research.

## Acknowledgments

This work was funded by public health service grants NIDA DA041480 (JRT), NIDA DA043443 (JRT), NIDA DA051598 (SMG), NIDA DA043533 (DJC) and McKnight Memory and Cognitive Disorders Award (DJC). Additional support was provided by the State of Connecticut through its support of the Ribicoff Laboratories. The views and opinions expressed in this manuscript are those of the authors and not shared by the State of Connecticut. The authors would like to acknowledge the useful discussions led by Matthew Roesch that made this collaborative work a possibility.

## References

Ahrens AM, Singer BF, Fitzpatrick CJ, Morrow JD, Robinson TE (2016) Rats that sign-track are resistant to Pavlovian but not instrumental extinction. Behav Brain Res 296:418–430 Available at: https://pubmed.ncbi.nlm.nih.gov/26235331/ [Accessed June 9, 2022].

Akam T, Rodrigues-Vaz I, Marcelo I, Zhang X, Pereira M, Oliveira RF, Dayan P, Costa RM (2021) The Anterior Cingulate Cortex Predicts Future States to Mediate Model-Based Action Selection. Neuron 109:149–163.e7 Available at: /pmc/articles/PMC7837117/ [Accessed May 17, 2021].

Amaya KA, Stott JJ, Smith KS (2020) Sign-tracking behavior is sensitive to outcome devaluation in a devaluation context-dependent manner: implications for analyzing habitual behavior. Learn Mem 27:136–149 Available at: https://pubmed.ncbi.nlm.nih.gov/32179656/ [Accessed June 9, 2022].

Boakes RA (1977) Performance on Learning to Associate a Stimulus with Positive Reinforcement. In: Operant-Pavlovian Interactions (Davis H, Hurwitz HMB, eds). ROUTLEDGE. Available at: https://www.routledge.com/Operant-Pavlovian-Interactions/Davis-Hurwitz/p/book/9780367713416 [Accessed March 21, 2022].

Chang SE, Todd TP, Bucci DJ, Smith KS (2015) Chemogenetic manipulation of ventral pallidal neurons impairs acquisition of sign-tracking in rats. Eur J Neurosci 42:3105–3116 Available at: https://onlinelibrary.wiley.com/doi/full/10.1111/ejn.13103 [Accessed February 8, 2022].

Cinotti F, Marchand AR, Roesch MR, Girard B, Khamassi M (2019) Impacts of inter-trial interval duration on a computational model of sign-tracking vs. goal-tracking behaviour. Psychopharmacology (Berl) 236:2373–2388 Available at: https://pubmed.ncbi.nlm.nih.gov/31367850/ [Accessed December 13, 2021].

Culbreth AJ, Westbrook A, Daw ND, Botvinick M, Barch DM (2016) Reduced model-based decision-making in schizophrenia. J Abnorm Psychol 125:777–787 Available at: http://doi.apa.org/getdoi.cfm?doi=10.1037/abn0000164 [Accessed February 22, 2018].

Daw ND, Gershman SJ, Seymour B, Dayan P, Dolan RJ (2011) Model-based influences on humans’ choices and striatal prediction errors. Neuron 69:1204–1215 Available at: http://www.ncbi.nlm.nih.gov/pubmed/21435563.

Dayan P, Berridge KC (2014) Model-based and model-free Pavlovian reward learning: Revaluation, revision, and revelation. Cogn Affect Behav Neurosci 14:473–492.

Derman RC, Schneider K, Juarez S, Delamater AR (2018) Sign-tracking is an expectancy-mediated behavior that relies on prediction error mechanisms. Learn Mem 25:550–563 Available at: http://learnmem.cshlp.org/content/25/10/550.full [Accessed June 1, 2022].

Deserno L, Huys QJM, Boehme R, Buchert R, Heinze H-J, Grace AA, Dolan RJ, Heinz A, Schlagenhauf F (2015) Ventral striatal dopamine reflects behavioral and neural signatures of model-based control during sequential decision making. Proc Natl Acad Sci U S A 112:1595–1600 Available at: http://www.ncbi.nlm.nih.gov/pubmed/25605941 [Accessed January 27, 2015].

Doñamayor N, Ebrahimi C, Garbusow M, Wedemeyer F, Schlagenhauf F, Heinz A (2021) Instrumental and Pavlovian Mechanisms in Alcohol Use Disorder. Curr Addict Reports 8:156–180 Available at: https://link.springer.com/article/10.1007/s40429-020-00333-9 [Accessed June 9, 2022].

Everett NA, Carey HA, Cornish JL, Baracz SJ (2020) Sign tracking predicts cue-induced but not drug-primed reinstatement to methamphetamine seeking in rats: Effects of oxytocin treatment. J Psychopharmacol 34:1271–1279 Available at: https://pubmed.ncbi.nlm.nih.gov/33081558/ [Accessed June 1, 2022].

Fitzpatrick CJ, Geary T, Creeden JF, Morrow JD (2019) Sign-tracking behavior is difficult to extinguish and resistant to multiple cognitive enhancers. Neurobiol Learn Mem 163 Available at: https://pubmed.ncbi.nlm.nih.gov/31319166/ [Accessed March 21, 2022].

Fitzpatrick CJ, Gopalakrishnan S, Cogan ES, Yager LM, Meyer PJ, Lovic V, Saunders BT, Parker CC, Gonzales NM, Aryee E, Flagel SB, Palmer AA, Robinson TE, Morrow JD (2013) Variation in the Form of Pavlovian Conditioned Approach Behavior among Outbred Male Sprague-Dawley Rats from Different Vendors and Colonies: Sign-Tracking vs. Goal-Tracking. PLoS One 8:e75042 Available at: https://journals.plos.org/plosone/article?id=10.1371/journal.pone.0075042 [Accessed December 9, 2021].

Flagel SB, Akil H, Robinson TE (2009) Individual differences in the attribution of incentive salience to reward-related cues: Implications for addiction. Neuropharmacology 56:139–148.

Flagel SB, Clark JJ, Robinson TE, Mayo L, Czuj A, Willuhn I, Akers CA, Clinton SM, Phillips PEM, Akil H (2011) A selective role for dopamine in stimulus-reward learning. Nature 469:53–59 Available at: https://pubmed.ncbi.nlm.nih.gov/21150898/ [Accessed December 13, 2021].

Flagel SB, Watson SJ, Akil H, Robinson TE (2008) Individual differences in the attribution of incentive salience to a reward-related cue: influence on cocaine sensitization. Behav Brain Res 186:48–56 Available at: https://pubmed.ncbi.nlm.nih.gov/17719099/ [Accessed June 1, 2022].

Groman S, Lee D, Taylor JR (2021) Unlocking the reinforcement-learning circuits of the orbitofrontal cortex. Behav Neurosci 135:120–128 Available at: https://pubmed.ncbi.nlm.nih.gov/34060870/ [Accessed July 23, 2021].

Groman SM, Massi B, Mathias SR, Curry DW, Lee D, Taylor JR (2019a) Neurochemical and behavioral dissections of decision-making in a rodent multistage task. J Neurosci 39.

Groman SM, Massi B, Mathias SR, Lee D, Taylor JR (2019b) Model-Free and Model-Based Influences in Addiction-Related Behaviors. Biol Psychiatry 85:936–945 Available at: https://linkinghub.elsevier.com/retrieve/pii/S0006322318321218 [Accessed September 21, 2019].

Groman SM, Thompson SL, Lee D, Taylor JR (2022) Reinforcement learning detuned in addiction: integrative and translational approaches. Trends Neurosci 45:96–105.

Hammersley R (1992) Cue exposure and learning theory. Addict Behav 17:297–300 Available at: https://pubmed.ncbi.nlm.nih.gov/1353283/ [Accessed December 13, 2021].

Hearst E, Jenkins HM (1974) Sign-trackingL: the stimulus-reinforcer relation and directed action. Austin Tex.: Psychonomic Society.

Hollerman JR, Schultz W (1998) Dopamine neurons report an error in the temporal prediction of reward during learning. Nat Neurosci 1:304–309 Available at: http://www.ncbi.nlm.nih.gov/entrez/query.fcgi?cmd=Retrieve&db=PubMed&dopt=Citation&list_uids=10195164.

Huys QJM, Tobler PN, Hasler G, Flagel SB (2014) The role of learning-related dopamine signals in addiction vulnerability 3. Prog Brain Res 211 Available at: http://dx.doi.org/10.1016/B978-0-444-63425-2.00003-9 [Accessed September 13, 2018].

Kawa AB, Bentzley BS, Robinson TE (2016) Less is more: prolonged intermittent access cocaine self-administration produces incentive-sensitization and addiction-like behavior. Psychopharmacology (Berl) 233:3587–3602 Available at: https://pubmed.ncbi.nlm.nih.gov/27481050/ [Accessed June 9, 2022].

Keefer SE, Bacharach SZ, Kochli DE, Chabot JM, Calu DJ (2020) Effects of Limited and Extended Pavlovian Training on Devaluation Sensitivity of Sign- and Goal-Tracking Rats. Front Behav Neurosci 14 Available at: https://pubmed.ncbi.nlm.nih.gov/32116587/ [Accessed December 13, 2021].

Keiflin R, Janak PH (2015) Dopamine Prediction Errors in Reward Learning and Addiction: From Theory to Neural Circuitry. Neuron 88:247–263 Available at: https://pubmed.ncbi.nlm.nih.gov/26494275/ [Accessed December 13, 2021].

Keiflin R, Pribut HJ, Shah NB, Janak PH (2019) Ventral Tegmental Dopamine Neurons Participate in Reward Identity Predictions. Curr Biol 29:93–103.e3 Available at: http://www.ncbi.nlm.nih.gov/pubmed/30581025 [Accessed February 5, 2020].

Kuhn BN, Campus P, Flagel SB (2018) The Neurobiological Mechanisms Underlying Sign-Tracking Behavior. In: Sign-Tracking and Drug Addiction (Tomie A, Morrow J, eds). Michigan Publishing, University of Michigan Library.

Lee B, Gentry RN, Bissonette GB, Herman RJ, Mallon JJ, Bryden DW, Calu DJ, Schoenbaum G, Coutureau E, Marchand AR, Khamassi M, Roesch MR (2018) Manipulating the revision of reward value during the intertrial interval increases sign tracking and dopamine release. PLOS Biol 16:e2004015 Available at: https://journals.plos.org/plosbiology/article?id=10.1371/journal.pbio.2004015 [Accessed March 21, 2022].

Lesaint F, Sigaud O, Flagel SB, Robinson TE, Khamassi M (2014a) Modelling Individual Differences in the Form of Pavlovian Conditioned Approach Responses: A Dual Learning Systems Approach with Factored Representations. PLoS Comput Biol 10:e1003466.

Lesaint F, Sigaud O, Flagel SB, Robinson TE, Khamassi M (2014b) Modelling individual differences in the form of Pavlovian conditioned approach responses: a dual learning systems approach with factored representations. PLoS Comput Biol 10 Available at: https://pubmed.ncbi.nlm.nih.gov/24550719/ [Accessed December 13, 2021].

Miller KJ, Botvinick MM, Brody CD (2017) Dorsal hippocampus contributes to model-based planning. Nat Neurosci 20:1269–1276 Available at: http://www.nature.com/doifinder/10.1038/nn.4613 [Accessed February 22, 2018].

Morrison SE, Bamkole MA, Nicola SM (2015) Sign tracking, but not goal tracking, is resistant to outcome devaluation. Front Neurosci 9:468.

Nasser HM, Calu DJ, Schoenbaum G, Sharpe MJ (2017) The Dopamine Prediction Error: Contributions to Associative Models of Reward Learning. Front Psychol 8 Available at: https://pubmed.ncbi.nlm.nih.gov/28275359/ [Accessed December 13, 2021].

Nasser HM, Chen YW, Fiscella K, Calu DJ (2015) Individual variability in behavioral flexibility predicts sign-tracking tendency. Front Behav Neurosci 9 Available at: https://pubmed.ncbi.nlm.nih.gov/26578917/ [Accessed December 13, 2021].

Pohořalá V, Enkel T, Bartsch D, Spanagel R, Bernardi RE (2021) Sign- and goal-tracking score does not correlate with addiction-like behavior following prolonged cocaine self-administration. Psychopharmacology (Berl) 238:2335–2346 Available at: https://pubmed.ncbi.nlm.nih.gov/33950271/ [Accessed June 1, 2022].

Robinson MJF, Berridge KC (2013) Instant transformation of learned repulsion into motivational “wanting.” Curr Biol 23:282–289 Available at: https://pubmed.ncbi.nlm.nih.gov/23375893/ [Accessed December 13, 2021].

Rode AN, Moghaddam B, Morrison SE (2020) Increased Goal Tracking in Adolescent Rats Is Goal-Directed and Not Habit-Like. Front Behav Neurosci 13 Available at: https://pubmed.ncbi.nlm.nih.gov/31992975/ [Accessed March 21, 2022].

Sadacca BF, Wikenheiser AM, Schoenbaum G (2017) Toward a theoretical role for tonic norepinephrine in the orbitofrontal cortex in facilitating flexible learning. Neuroscience 345:124–129 Available at: https://www.sciencedirect.com/science/article/pii/S0306452216300823 [Accessed September 9, 2019].

Saunders BT, Richard JM, Margolis EB, Janak PH (2018) Dopamine neurons create Pavlovian conditioned stimuli with circuit-defined motivational properties. Nat Neurosci 21:1072–1083.

Saunders BT, Robinson TE (2010) A Cocaine Cue Acts as an Incentive Stimulus in Some but not Others: Implications for Addiction. Biol Psychiatry 67:730–736.

Saunders BT, Robinson TE (2011) Individual variation in the motivational properties of cocaine. Neuropsychopharmacology 36:1668–1676.

Saunders BT, Robinson TE (2012) The role of dopamine in the accumbens core in the expression of Pavlovian-conditioned responses. Eur J Neurosci 36:2521–2532 Available at: https://pubmed.ncbi.nlm.nih.gov/22780554/ [Accessed December 13, 2021].

Saunders BT, Robinson TE (2013) Individual variation in resisting temptation: Implications for addiction. Neurosci Biobehav Rev 37:1955–1975.

Saunders BT, Yager LM, Robinson TE (2013) Cue-evoked cocaine “craving”: role of dopamine in the accumbens core. J Neurosci 33:13989–14000 Available at: https://pubmed.ncbi.nlm.nih.gov/23986236/ [Accessed June 9, 2022].

Sebold M, Schad DJ, Nebe S, Garbusow M, Jünger E, Kroemer NB, Kathmann N, Zimmermann US, Smolka MN, Rapp MA, Heinz A, Huys QJM (2016) Don’t Think, Just Feel the Music: Individuals with Strong Pavlovian-to-Instrumental Transfer Effects Rely Less on Model-based Reinforcement Learning. J Cogn Neurosci 28:985–995 Available at: http://direct.mit.edu/jocn/article-pdf/28/7/985/1951515/jocn_a_00945.pdf [Accessed December 12, 2021].

Sharpe MJ, Chang CY, Liu MA, Batchelor HM, Mueller LE, Jones JL, Niv Y, Schoenbaum G (2017) Dopamine transients are sufficient and necessary for acquisition of model-based associations. Nat Neurosci 20:735–742 Available at: http://www.ncbi.nlm.nih.gov/pubmed/28368385 [Accessed February 22, 2018].

Smedley EB, Smith KS (2018) Evidence for a shared representation of sequential cues that engage sign-tracking. Behav Processes 157:489–494 Available at: https://pubmed.ncbi.nlm.nih.gov/29933057/ [Accessed June 9, 2022].

Swintosky M, Brennan JT, Koziel C, Paulus JP, Morrison SE (2021) Sign tracking predicts suboptimal behavior in a rodent gambling task. Psychopharmacology (Berl) 238:2645–2660 Available at: https://pubmed.ncbi.nlm.nih.gov/34191111/ [Accessed December 13, 2021].

Tunstall BJ, Kearns DN (2015) Sign-tracking predicts increased choice of cocaine over food in rats. Behav Brain Res 281:222–228.

Uslaner JM, Acerbo MJ, Jones SA, Robinson TE (2006) The attribution of incentive salience to a stimulus that signals an intravenous injection of cocaine. Behav Brain Res 169:320–324 Available at: https://pubmed.ncbi.nlm.nih.gov/16527365/ [Accessed December 13, 2021].

Wang F, Schoenbaum G, Kahnt T (2020) Interactions between human orbitofrontal cortex and hippocampus support model-based inference. PLoS Biol 18 Available at: https://pubmed.ncbi.nlm.nih.gov/31961854/ [Accessed December 13, 2021].

